# Genome sequence data and integrative RNA-Seq data of Black Soldier Fly (*Hermetia illucens*) (A Japanese strain)

**DOI:** 10.1101/2025.07.08.663698

**Authors:** Koji Takeda, Takuya Uehara, Akiya Jouraku, Chia-Ming Liu, Tetsuya Kobayashi, Masami Shimoda, Kakeru Yokoi

**Author notes:** Corresponding author: Kakeru Yokoi (,). These authors contributed equally (Koji Takeda, Takuya Uehara, Akiya Jouraku).

## Abstract

The black soldier fly, *Hermetia illucens*, is a dipteran insect with a distinctive black body in the adult stage. The larvae of this species possess biological features which can be used for industrial applications. We established an *H. illucens* field-collected strain to develop an industrial system for recycling organic waste. We constructed whole-genome sequences of the strain of *H. illucens* and prepared a gene set and its functional annotation data to promote research on the social implementation of *H. illucens* and genetically characterise this strain. In addition, time-course transcriptome data of the larvae and transcriptome expression data of multiple tissues from 6th instar larvae and male and female adults were prepared. The genome size and its N50 value were approximately 1.05 Gbp and 191 bp, respectively. The results of the validation analyses of the genome and transcriptome expression data showed that these data were reliable as reference data, suggesting that they can be used in research on the industrial usage of evolutionary *H. illucens* or evolutionary or comparative biology.

## Background

The black soldier fly (BSF) (*Hermetia illucens*) is a dipteran insect with a distinctive black body colour in its adult stage, measuring 15–20 mm in length, with partly hyaline areas on the first and second abdominal segments. Black soldier fly larvae (BSFL) are 10–20 mm long, cylindrical, pale yellowish to brown in colour, and have a robust body structure^1^. In contrast to adults, who consume only minimal amounts of water and sugar dew^2^, BSFL efficiently decompose decaying organic matter and exhibit rapid growth and high feeding efficiency.

Their saprophagous nature enables them to feed and grow on various organic materials, such as food residues, animal manure, and even low-nutrient substrates such as coffee husks. This remarkable adaptability is one of the most distinct features of this species. Photographs of typical BSF at all developmental stages are shown in Fig. 1.

**Fig. 1.**
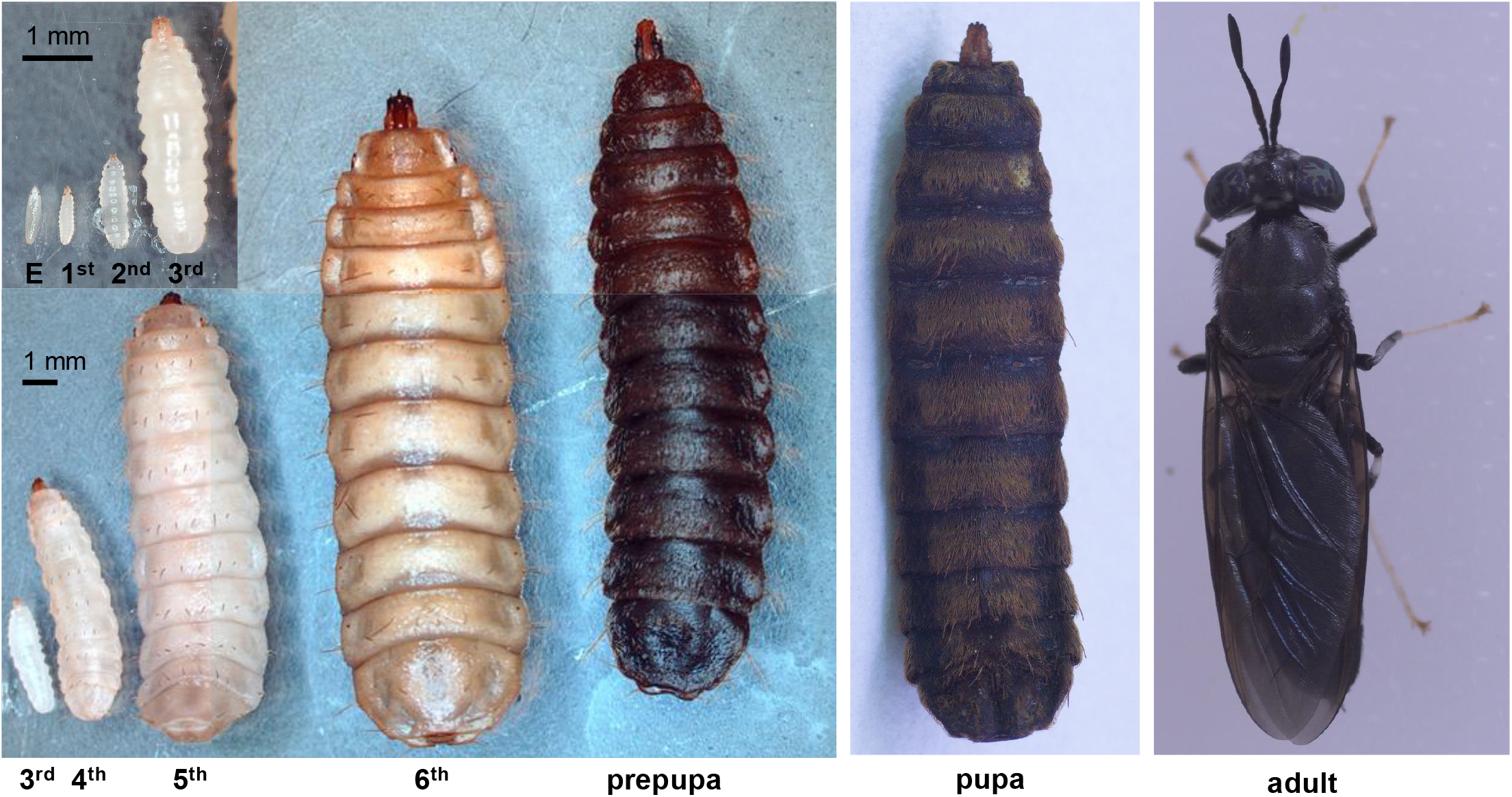
Photographs of the black soldier fly. Photographs of black soldier flies at each developmental stage: egg, 1st to 6th instar larvae, prepupa, pupa, and adult.

The biological features of BSFL have been exploited industrially for the efficient bioconversion of organic waste. For example, they can accumulate more than 40% of their body weight as lipids during the 14–18 day larval period^2,3^. These lipids are particularly rich in lauric acid (C12:0), which makes BSFL a promising resource for animal feed and biofuel production^4^. Additionally, BSFL are regarded as a high-quality source of protein. Defatted and dried BSFL meal can contain up to 42% crude protein^5^, drawing growing attention for its potential use in aquaculture and livestock feed^6^. Given the environmental challenges posed by conventional protein sources, such as fishmeal and soybeans, BSFL represents a sustainable and scalable alternative that can help address the global protein shortage. Therefore, BSFL is a valuable resource that contributes to the development of a recycling-oriented society.

BSF is believed to have originated in the Neotropical region, where its ancestral genetic diversity is concentrated^7^. Based on these genetic hotspots, this species has expanded its range over ancient times, eventually establishing populations on other continents. This global spread was driven primarily by human-mediated introductions, and BSF established its present-day distribution by the 1960s^8^. Despite the recent establishment of their current distribution, BSF harbours considerable hidden genetic diversity. Genomic studies have revealed the existence of at least two major genetic lineages that diverged more than three million years ago^9^. Substantial intraspecific variation has been observed in mitochondrial sequences^10,11^. Nonetheless, most of the commercial and laboratory populations currently used are thought to have originated from a single North American strain. This genetic uniformity poses a significant challenge for the stable industrial utilisation of BSF. A narrow genetic base may increase vulnerability to disease outbreaks and reduce adaptability to environmental stresses. Genetically diverse populations are crucial for selective breeding programs aimed at improving productivity, enhancing specific nutritional profiles, and conferring disease resistance. A comprehensive understanding of BSF’s genetic diversity and the development of breeding strategies that leverage this diversity are essential for sustainable and efficient industrial applications.

We established a laboratory strain from field-collected individuals in Japan and have since been developing applications for organic waste recycling and the industrial use of BSF, including improving rearing conditions^12^ and enhancing feed quality^13^. Although genomic resources have been developed in other countries^14-16^, no independent genetic studies on the evolutionary history of BSF have been conducted in Japan, and no distinct evolutionary lineages have been identified within the country. In this study, we genetically characterised the Japanese population, assembled a high-quality reference genome, and generated tissue- and age-specific gene expression profiles to support future breeding efforts using genomic and genome-editing technologies (Fig. 2): RNA-Seq data were prepared for the time-course BSF larval stages (T1–T8 and P); the full gut (FG), midgut (MG), hindgut (HG), Malpighian tubules (MT), white Malpighian tubules (WMT), and fatbodies (FB) of last-day 6th instar larvae; and the whole body (Mw and Fw), head (Mh and Fh), and tarsus (Mt and Ft) of both male and female adults.

**Fig. 2.**
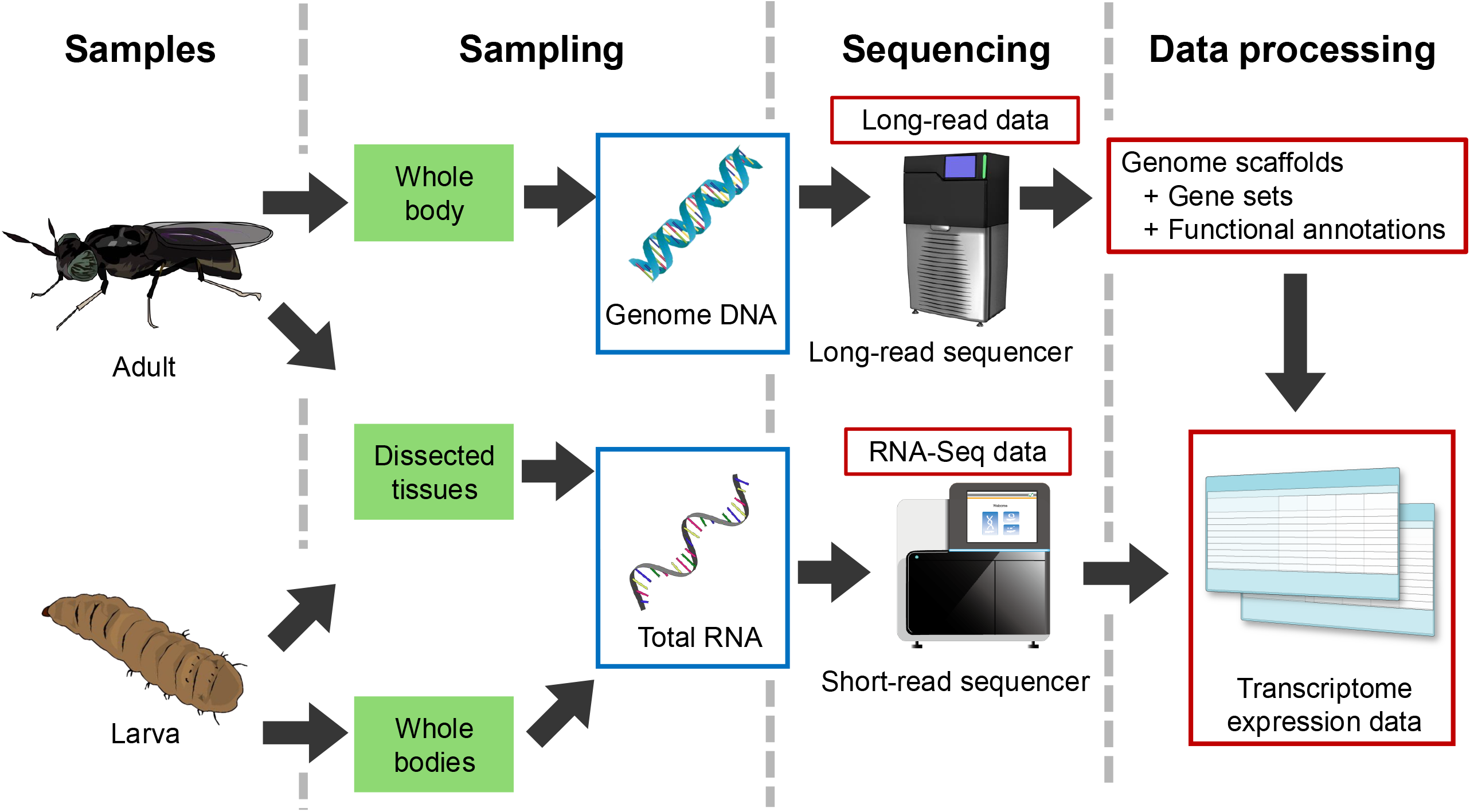
Workflow of this study. Schematic workflow of this study Blue boxes indicate genomic DNA or total RNA. Red boxes indicate sequence or related data.

## Methods

### Sample Preparation

#### Insect Rearing

Black soldier flies were originally collected from Tsukuba, Japan, in 2013. The rearing method was modified from Nakamura et al. (2016)^17^. Briefly, adult flies were housed in mesh cages (W 115 × D 85 × H 85 cm) and allowed to mate and oviposit. Eggs were collected using an oviposition device, and the oviposition date was recorded. The collected eggs were transferred to plastic cups and incubated at 28 °C until hatching. The day of hatching was designated day 1 and served as the starting point for all subsequent experiments. Newly hatched larvae were fed an artificial diet^13^ and reared in an incubator maintained at 28 °C.

#### DNA extraction and sequencing

DNA was extracted from a newly eclosed adult male fly. High-molecular-weight DNA was obtained using a gravity flow extraction kit (Genomic-tip 100/G, Cat. no. 10243; QIAGEN, Hilden, Germany). The extracted DNA was used for long-read sequencing. Samples were prepared following the SMRTbell preparation guide for the PacBio Sequel II System. Sequencing was performed by Macrogen Japan Corp (Tokyo, Japan) using SMRT technology.

#### RNA Extraction and sequencing

Whole-body samples (6th instar larvae, pupae, and adults; Table 3) were collected from single individuals, whereas specific tissues (head and tarsus) were dissected from three individuals and pooled. All samples were homogenised on ice using a plastic pestle, and RNA was extracted using TRIzol reagent according to the manufacturer’s protocol. Larval samples were collected at distinct developmental stages, with each sample comprising ten individuals. Larvae with recorded hatching dates were reared on an artificial diet and collected at two-day intervals. The body weights of the harvested larvae were measured, and ten individuals with comparable weights were selected for RNA extraction. The larvae were rinsed with tap water followed by ultrapure water, pooled into a single tube, and homogenised using a disposable plastic homogeniser (Nippi, Tokyo, Japan). Total RNA was extracted using an RNeasy Kit (QIAGEN) following the manufacturer’s instructions. Extracted RNA samples were stored at −80 °C until further processing. The sequencing library was prepared using random fragmentation of the DNA or cDNA sample, followed by 5′ and 3′ adapter ligation. Alternatively, “tagmentation” combines the fragmentation and ligation reactions into a single step, greatly increasing the efficiency of the library preparation process. The adapter-ligated fragments were then PCR-amplified and gel-purified. For cluster generation, the library was loaded into a flow cell, where fragments were captured on a lawn of surface-bound oligos complementary to the library adapters. Each fragment was amplified into distinct clonal clusters using bridge amplification. The templates were ready for sequencing using an Illumina NovaSeq 6000 after cluster generation was complete. Sequencing library construction and sequencing were performed by Macrogen Japan Corp.

#### Sequence data analysis

All scripts and commands (settings and options) of the data analysis in this study are available via the “Code Availability” section.

#### Genome sequence construction and gene prediction

*De novo* assembly was initially performed using CANU version 2.1.1^18^ using long-read sequencing data. The corrected reads generated by CANU were subsequently used as inputs for another *de novo* genome assembler using Hifiasm version 0.14-r312^19-21^ to improve genome continuity. The resulting contigs were polished with NextPolish version 1.3.1, scaffolded using ragtag version 2.1.0^22^, with reference to the published BSF genome^23^, resulting in the final genome assembly. The repeat regions in the genome were masked for gene model prediction using RepeatModeler version 2.0.2a and RepeatMasker version 4.1.2. pl^24,25^. RNA-Seq reads (L, P, Fh, Ft, Fw, Mh, Mt, and Mw) were aligned to the repeat-masked genome using STAR version 2.7.6a to generate hints for gene prediction. Gene prediction was performed using BRAKER version 2.1.6 based on both the mapped RNA-seq hints and *Drosophila melanogaster* protein data^26^. The two resulting gene predictions were merged into a unified gene prediction using TSEBRA version 1.0.3. Functional annotation of predicted gene sets was performed using various tools. Homologous protein searches by BLASTP version 2.12.0+ with option “-evalue 1e-3”^27^ were conducted for protein sequences of NCBI nr database (retrieved on 2023-10-23) and *D. melanogaster*^26^. Homologous domain searches were conducted by HMMER version 3.1b2^28^ using Pfam version 35.0^29^. Classification of proteins (identifying related InterPro entries and Gene Ontology (GO) terms) was conducted using InterProScan version 5.55-88.0^30^ with the options “-dp -iprlookup - goterms”.

#### RNA-seq data analysis

RNA-seq raw reads of each sample were cleaned using BBTools ver 39.06 (https://sourceforge.net/projects/bbmap/) with the bbduk.sh command^31^. Clean reads were mapped to the reference genome constructed in this study using STAR version 2.7.10b^32^. The genome indices of STAR were constructed using the “STAR” command. Mapping clean reads using the STAR genome indices was conducted using the “STAR” command. The expression values (expected count and TPM) of genes and transcripts in each sample were calculated using RSEM version 1.3.3^33^ using the mapping data. The transcriptome indices of RSEM were constructed using the “rsem-prepare-reference” command. Then, the expression values were calculated using the “rsem-calculate-expression” command.

Unmapped RNA-seq read pairs of the “T1” sample were extracted to narrow down the cause of the low mapping rate (58.61%) observed in “T1” sample by rerunning STAR with an additional option “--outReadsUnmapped Fastx”. Several unmapped read pairs were compared to the NCBI nt database using BLASTN version 2.12.0+. All top-hit sequences in the nt database were derived from mRNA of *Saccharomyces cerevisiae*. Therefore, the low mapping rate was attributed to the contamination of *S. cerevisiae*. All unmapped reads were mapped to the reference genome of *Saccharomyces cerevisiae strain* S288C^34^ using STAR in the same manner as described above to confirm the estimation.

#### Busco analysis and hierarchical clustering analysis

Busco analysis was performed using BUSCO version 5.2.2, with the insecta_odb10 database and the assembled genome sequence data as input, utilising the default settings^35^. Hierarchical clustering analysis was performed using R (version 4.2.3) in RStudio (version 2023.03.0+386), using transcriptome expression matrix data as input. The principal component analysis (PCA) was performed using R.

#### Data records

All sequencing data were deposited in the DNA Data Bank of Japan (DDBJ) Sequence Read Archive (SRA). The accession ID of the long-read DNA sequence was DRR680776, and the RNA-Seq IDs are listed in Table 1. The constructed genome sequence is available in DDBJ GenBank. The following are available in Figshare: a list of sequence contig names and accession IDs (DOI: 10.6084/m9.figshare.29422424), detailed output data related to RepeatModeler and RepeatMasker (DOI: 10.6084/m9.figshare.29423243), and the predicted gene set data and functional annotation result files (gtf, fasta, and xlsx) (DOI: 10.6084/m9.figshare.29423210). The calculated transcriptome expression value matrix files were deposited into the DDBJ Genomic Expression Archive (GEA) (Accession ID: E-GEAD-1083 and E-GEAD-1083), and merged files from the two expression matrix files used for hierarchical clustering analysis are available in Figshare (DOI: 10.6084/m9.figshare.29423267).

**Table 1.** RNA-Seq sample and data list.

| Sample name* | Sample name in SRA or GEA | Tissue | Developmental stage | Days after hatching | Sex | SRA run ID |
| --- | --- | --- | --- | --- | --- | --- |
| T1 | A-01 | Whole body | 1st-2nd instar larvae | 1 | - | DRR667142 |
| T2 | A-02 | Whole body | 3rd-4th instar larvae | 3 | - | DRR667143 |
| T3 | A-03 | Whole body | 5th instar larvae | 5 | - | DRR667144 |
| T4 | A-04 | Whole body | 5th instar larvae | 7 | - | DRR667145 |
| T5 | A-05 | Whole body | 5th instar larvae | 9 | - | DRR667146 |
| T6 | A-06 | Whole body | 6th instar larvae | 11 | - | DRR667147 |
| T7 | A-07 | Whole body | 6th instar larvae | 12 | - | DRR667148 |
| T8 | A-08 | Whole body | 6th instar larvae | 16 | - | DRR667149 |
| L | L-1 | Whole body | 6th instar larvae | - | - | DRR667134 |
| FG1 | FG1 | Full Gut | 6th instar larvae | - | - | DRR502194 |
| FG2 | FG2 | Full Gut | 6th instar larvae | - | - | DRR502195 |
| FG3 | FG3 | Full Gut | 6th instar larvae | - | - | DRR502196 |
| HG | B-11 | Hindgut | 6th instar larvae | 20 | - | DRR667150 |
| MG1 | MG1 | Midgut | 6th instar larvae | - | - | DRR502197 |
| MG2 | MG2 | Midgut | 6th instar larvae | - | - | DRR502198 |
| MG3 | MG3 | Midgut | 6th instar larvae | - | - | DRR502199 |
| MG4 | B-12 | Midgut | 6th instar larvae | 20 | - | DRR667151 |
| MT1 | MT1 | Malpighian tubules | 6th instar larvae | - | - | DRR502200 |
| MT2 | MT2 | Malpighian tubules | 6th instar larvae | - | - | DRR502201 |
| MT3 | B-13 | Malpighian tubules | 6th instar larvae | 20 | - | DRR667152 |
| WMT1 | WT1 | White Malpighian tubules | 6th instar larvae | - | - | DRR502202 |
| WMT2 | WT2 | White Malpighian tubules | 6th instar larvae | - | - | DRR502203 |
| WMT3 | B-14 | White Malpighian tubules | 6th instar larvae | 20 | - | DRR667153 |
| FB | B-15 | Fatbody | 6th instar larvae | 20 | - | DRR667154 |
| P | P-1 | Pupa | Pupa | - | - | DRR667141 |
| Fh1 | fh-1 | Head | Adult | - | Female | DRR667128 |
| Fh2 | fh-2 | Head | Adult | - | Female | DRR667129 |
| Ft1 | ft-1 | Tarsus | Adult | - | Female | DRR667130 |
| Ft2 | ft-2 | Tarsus | Adult | - | Female | DRR667131 |
| Fw1 | fw-1 | Whole body | Adult | - | Female | DRR667132 |
| Fw2 | fw-2 | Whole body | Adult | - | Female | DRR667133 |
| Mh1 | mh-1 | Head | Adult | - | Male | DRR667135 |
| Mh2 | mh-2 | Head | Adult | - | Male | DRR667136 |
| Mt1 | mt-1 | Tarsus | Adult | - | Male | DRR667137 |
| Mt2 | mt-2 | Tarsus | Adult | - | Male | DRR667138 |
| Mw1 | mw-1 | Whole body | Adult | - | Male | DRR667139 |
| Mw2 | mw-2 | Whole body | Adult | - | Male | DRR667140 |

\*Numbers added after sample name indicate biological replicate

## Results and Discussion

### Genome sequence, gene set data and Transcriptome data

The size and N50 value of the constructed genome were approximately 1.05 Gbp and 191 Mbp, respectively. The genome data consisted of 66 contigs, of which the average length and largest contig size were approximately 15 and 234 Mbp, respectively (Table 2). The proportion of CG sequences was 42.36%. The repetitive regions and transposable elements (consisting of multiple types of the elements) comprised 68% of the genome sequence regions (Table 3). Using the genome sequence data, 17,892 genes and 18,830 transcripts were predicted (gene set data) (Table 2), and these genes were functionally annotated (functional annotation data) (Fig. 2). A summary of the functional annotations is presented in Table 4. As described above, the transcriptome expression matrix data of time-course larval stages and multiple tissues of larvae and adults were prepared, which included the time-course BSF larval stages (T1–T8 and P); FG, MG, HG, MT, WMT, and FB of last-day 6th instar larvae; and the whole body (Mw and Fw), head (Mh and Fh), and tarsus (Mt and Ft) of both male and female adults (Fig. 2 and Table 1).

**Table 2.**
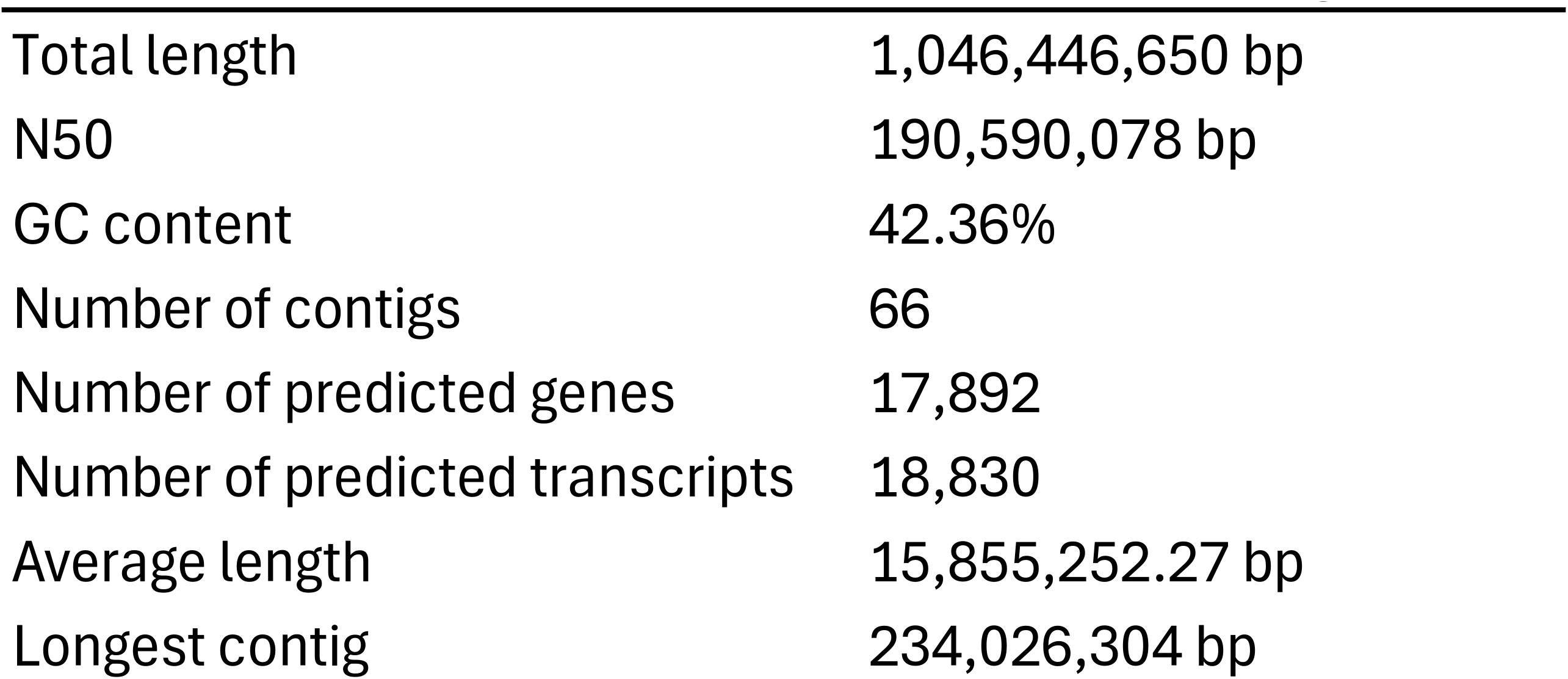
Basic statistics of the constructed *H. illucens genome*.

**Table 3.** Repetitive regions and transposable elements in the assembled BSF genome.

|  | Number of elements* | Length oppcupied | Percentage |
| --- | --- | --- | --- |
| SINEs | 23,294 | 4,700,919 bp | 0.45% |
| LINEs | 1,034,728 | 459,705,977 bp | 43.93% |
| LINE2 | 30,083 | 12,883,029 bp | 1.23% |
| L3/CR1 | 24,329 | 14,248,601 bp | 1.36% |
| LTR elements | 83,219 | 43,817,987 bp | 4.19% |
| DNA elements | 124,789 | 42,311,076 bp | 4.04% |
| Unclassified | 564,027 | 142,093,847 bp | 13.58% |
| **Total interspersed repeats** |  | **692,629,806 bp** | **66.19%** |
| Small RNA | 24,472 | 4,809,580 bp | 0.46% |
| Simple repeats: | 53,193 | 4,150,315 bp | 0.40% |
| Low complexity | 7713 | 370,992 bp | 0.04% |
| **Total TE and repetetive region** |  |  | **67%** |

\* most repeats fragmented by insertions or deletions have been counted as one element

**Table 4.** Summary of the functional annotation for the predicted gene set (17,892 genes)

| Analysis Tool | Database | No. of annotated genes | Annotation rate (%) |
| --- | --- | --- | --- |
| BLASTP | NCBI nr (2023-10-23) | 17,342 | 96.92 |
| BLASTP | *Drosophila melanogaster* (GCF_000001215.4) | 13,428 | 75.05 |
| HMMER | Pfam 35.0 | 14,006 | 78.28 |
| InterProScan | InterPro entries in InterProScan 5.55-88.0 | 13,600 | 76.01 |
| InterProScan | GO terms in InterProScan 5.55-88.0 | 9,763 | 54.57 |
| Total |  | 17,403 | 97.27 |

### Genome sequence data validation

As shown in Table 2, the constructed genome size was approximately 1.05 Gbp, which is comparable to those of the two published BSF genomes (1.1 Gbp and 1.01 Gbp)^14,16^ while the N50 value was higher than those of the two genome datasets. The number of predicted genes (17,892) did not differ significantly from that of a published gene set (16,770 genes)^14^. In addition, BUSCO analysis was performed to assess the quality of the constructed BSF genome data (Table 5). The results showed that ratio of Complete BUSCO was 98.7%, comparable to the BUSCO results of the published other BSF genome data^14,16^. This indicated that the constructed genome data possessed sufficient quality to serve as reference genome data.

**Table 5.** Busco result (Using insecta_odb10, total number of BUSCOs: 1367)

| **BUSCO group** | **Number of**<br>**found**<br>**BUSCOs**<br>**(percent of**<br>**all BUSCOs)** |
| --- | --- |
| Complete BUSCOs (C) | 1349 (98.7%) |
| Complete and single-copy BUSCOs (S) | 1290 (98.7%) |
| Complete and duplicated BUSCOs (D) | 59 (4.3%) |
| Missing BUSCOs (M) | 13 (0.9%) |
| Fragmented BUSCOs (F) | 5 (0.4%) |
| Total BUSCOs | 1367 |

### RNA-seq data validation

The mapping rates of RNA-seq samples ranged from 93 to 98% (Table 6), indicating high mapping rates for the genome assembly. However, exceptionally, the mapping rate of the “T1” sample was 58.61%. Since 88.33% of the unmapped reads in the “T1” sample were mapped to the *S. cerevisiae* genome assembly, the cause of the low mapping rate was confirmed due to the contamination of *S. cerevisiae*. Although the mapping rate of the “T1” sample was lower than that of other samples, the expression values of the “T1” sample calculated using their mapped reads can still be useful for comparison with other samples. A PCA analysis of the dataset (RNA samples of T1−T8, HG, MT3, MG4, WMT3 and FB sequenced at the same time) was conducted using the plotPCA function in DESeq2 ver 1.44.0 (Love et al. 2014) to evaluate the validity of the “T1” sample. The expression values (expected counts of all genes) were normalized among the samples using the vst function in DESeq2 ver 1.44.0 (genes with low total expected counts (<10) in all the samples were discarded for the analysis). The results of PCA analysis showed that PC1 and PC2 explained 65% and 11% of the variance, respectively (Fig. 3). The normalized expression values of the “T1” sample were considered suitable for comparison with other samples because the whole-body samples from different larval stages (“T1”–”T8”) formed a distinct cluster. Additionally, the “T1” sample (0 days) was positioned relatively close to the “T2” sample (2 days), compared to the later-stage samples (4–16 days).

**Table 6.** RNA-seq sample statistics.

| Sample ID | No. of raw reads | No. of clean reads | Mapping rate (%) |
| --- | --- | --- | --- |
| T1 | 63,729,994 | 60,755,908 | 58.61 |
| T2 | 41,156,590 | 39,466,784 | 97.8 |
| T3 | 60,949,934 | 58,230,654 | 97.73 |
| T4 | 55,665,680 | 53,044,804 | 97.63 |
| T5 | 41,442,092 | 39,324,586 | 97.7 |
| T6 | 59,675,144 | 56,289,346 | 97.72 |
| T7 | 18,572,118 | 17,478,750 | 98.49 |
| T8 | 48,708,950 | 45,743,072 | 97.49 |
| L | 53,824,580 | 50,401,028 | 96.45 |
| FG1 | 25,932,084 | 24,749,212 | 96.32 |
| FG2 | 22,308,952 | 21,227,348 | 95.02 |
| FG3 | 26,192,044 | 24,970,358 | 95.38 |
| HG | 42,655,354 | 40,808,394 | 97.48 |
| MG1 | 20,066,512 | 19,128,064 | 97.36 |
| MG2 | 27,409,840 | 26,022,446 | 96.71 |
| MG3 | 23,538,962 | 22,440,922 | 96.78 |
| MG4 | 58,465,366 | 55,957,938 | 97.72 |
| MT1 | 26,502,612 | 25,191,510 | 95.37 |
| MT2 | 26,218,224 | 24,830,460 | 95.11 |
| MT3 | 41,279,572 | 39,211,114 | 95.59 |
| WMT1 | 27,443,392 | 25,924,338 | 96.14 |
| WMT2 | 25,353,956 | 24,067,096 | 96.65 |
| WMT3 | 64,523,388 | 61,480,364 | 95.76 |
| FB | 55,842,144 | 53,355,722 | 98 |
| P | 42,762,176 | 39,658,022 | 95.57 |
| Fh1 | 44,531,784 | 41,177,262 | 93.42 |
| Fh2 | 40,303,404 | 37,369,622 | 93.48 |
| Ft1 | 48,375,608 | 45,437,064 | 95.95 |
| Ft2 | 43,845,062 | 40,884,322 | 95.76 |
| Fw1 | 52,686,736 | 49,243,664 | 95.84 |
| Fw2 | 54,095,328 | 50,568,392 | 95.59 |
| Mh1 | 52,304,926 | 48,500,902 | 93.97 |
| Mh2 | 44,191,108 | 41,142,496 | 93.69 |
| Mt1 | 50,471,152 | 47,072,344 | 95.58 |
| Mt2 | 44,464,020 | 41,218,664 | 95.52 |
| Mw1 | 40,675,700 | 37,682,222 | 95.38 |
| Mw2 | 52,490,950 | 49,152,946 | 95.65 |

**Fig. 3.**
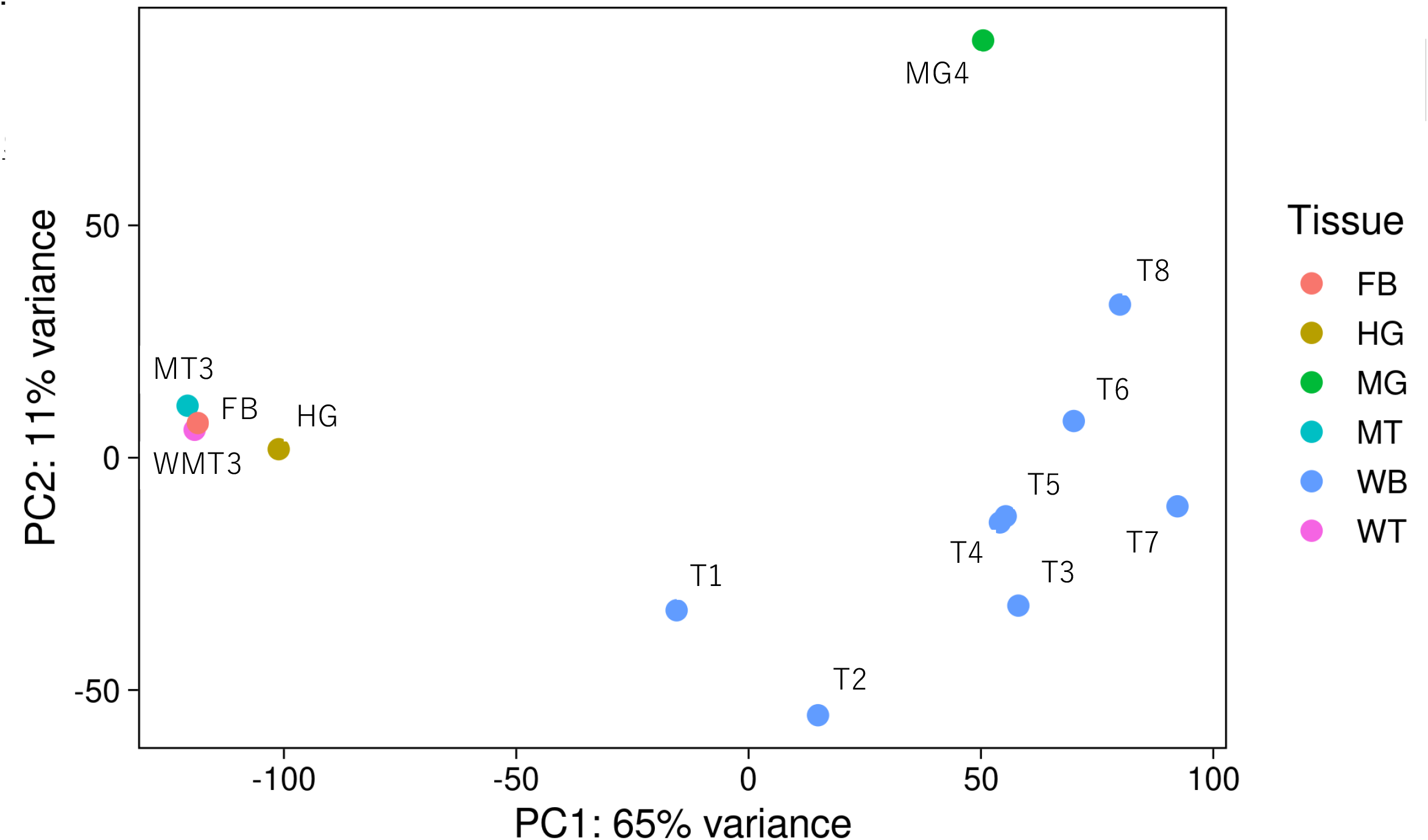
Principal component analysis (PCA) results of expression data of certain samples. The horizontal and vertical axes are the first and second principal components, respectively. Blue dots indicate time-course larvae samples (WB). Dots representing other tissue samples are shown in various colours, as indicated in the legend on the right side of the PCA plot.

Hierarchical clustering analysis using all transcriptome expression data was performed (Fig. 4) to assess the reliability of transcriptome expression data. Adult, pupal, and larval samples were separated into large left and right clusters, respectively. Time-course larval samples (T1–T8), except T7, formed one cluster, whereas L and T7 samples were placed at different locations from the clusters. This may be because samples L and T7 were in the pupation preparation phase. The MG and FG samples formed independent clusters. WMT, MT, and FB samples formed one cluster, which may be because WMT, MT, and FB are spatially close to each other in the larval body. The Fh and Mh samples formed one cluster, whereas the Ft and Mt samples formed one cluster. In addition, Fw and P formed a single cluster. The Mw samples were separated from those of the other adults. This may be because the Mw samples contained testes, in which expression data are different from those of other tissues in the silkworm^36^. These results indicated that the clustering results were mostly reasonable, suggesting that the RNA-Seq and expression data are reliable as reference expression data.

**Fig. 4.**
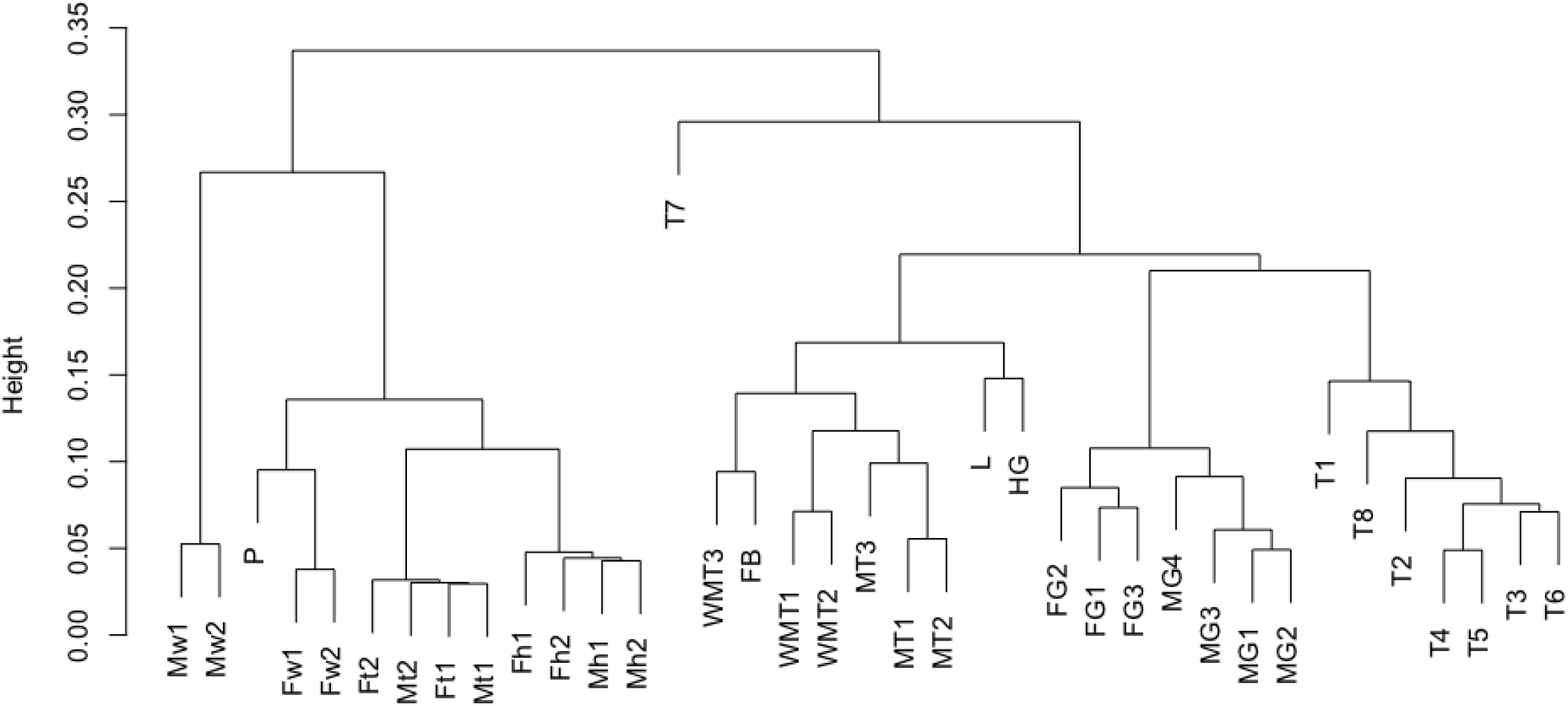
Dendrogram of hierarchical clustering analysis using transcriptome expression data. The sample names in the dendrogram and metadata are shown in Table 3.

As described in the “Technical Validation” section, the constructed genome data and RNA-Seq data were reliable as reference genome and transcriptome data. These data contribute to evolutionary and comparative biology. Furthermore, using these data, genome editing target genes can be determined, which will lead to the breeding of strains possessing beneficial phenotypes for the commercial usage of BSF as food or feed.

## Code Availability

All script codes and commands for data analysis in this study are available in figshare (DOI: 10.6084/m9.figshare.29423852).

## Acknowledgements

This work was supported by the Cabinet Office, Government of Japan, Cross-ministerial Moonshot Agriculture, Forestry and Fisheries Research and Development Program, and “Technologies for Smart Bio-industry and Agriculture” (funding agency: Bio-oriented Technology Research Advancement Institution) [JPJ009237]. Parts of Fig. 2 were drawn using illustrations from TogoTV (© 2016 DBCLS TogoTV, CC-BY-4.0 https://creativecommons.org/licenses/by/4.0/deed.ja).

We would like to thank Editage (www.editage.jp) for English language editing.

## Author contributions

K.T., T.U., C.M.L, T.K. and M.S. conceived the study. K.T., T.U., C.M.L, T.K. and M.S. prepared the genome and RNA samples. T.U., A.J., and K.Y. performed bioinformatics data analysis. K.T., T.U., C.M.L, A.J. and K.Y wrote the original draft of the manuscript. All authors reviewed and edited the draft of the manuscript. All the authors have read and agreed to the published version of the manuscript.

